# Pmt1-dependent and state-restricted DNA methylation in fission yeast

**DOI:** 10.1101/2025.09.01.672383

**Authors:** Samia Miled

## Abstract

DNA 5-methylcytosine (5mC) is widely considered absent from budding and fission yeasts. The Dnmt2 family enzyme Pmt1 in *Schizosaccharomyces pombe* is annotated as a tRNA C38 methyltransferase, but whether it methylates DNA in vivo has remained unresolved. Using orthogonal chemistry (HPLC/LC-MS and 1H-NMR) with external standards (limit of detection for 5mdC = 0.0125 mM), we detect authentic 5-methyl-2’- deoxycytidine (5mdC) in enzymatically digested genomic DNA from *S. pombe*. Quantification across isogenic strains shows that 5mdC is undetectable in vegetative wild type but rises to 0.518% (plus or minus 0.025) of cytosines two hours after G0 exit. Loss of the cytosine/5-mC deaminase Fcy1 causes marked accumulation: 1.445% (plus or minus 0.361) in vegetative cells and 5.943% (plus or minus 1.364) at 2 hours, approximately 11.5-fold over wild type. By contrast, *pmt1Δ* and *fcy1Δ pmt1Δ* remain below detection in all conditions, establishing Pmt1 as the DNA-directed methyltransferase in vivo. Nascent-strand fractionation and lambda exonuclease enrichment place this transient 5mdC pulse on Okazaki-enriched DNA, consistent with co-replicative installation. Functionally, the first S phase after quiescence shows increased Rad22-YFP foci in *fcy1Δ*, which are dampened by queuine ; in a sensitized background (*ung1Δ thp1Δ*), deleting *fcy1* reduces C->T transitions by about 30 percent. Together, chemical, genetic, and temporal evidence reveals a Pmt1- dependent, state-restricted DNA methylation program in fission yeast that is rapidly curtailed by Fcy1, redefining the epigenetic landscape of S. pombe and providing a minimal, tractable system to dissect regulated DNA methylation in eukaryotes.

## Introduction

Cytosine 5-methylation (5mC) is a pervasive epigenetic mark across eukaryotes, but its distribution is highly uneven across lineages. Early comparative surveys established a canonical view in which *Schizosaccharomyces pombe* lacks detectable genomic 5mC, in stark contrast to many plants and animals [1–3]. The strength of that consensus has rested largely on bisulfite-based assays and limited mass-spectrometric analyses conducted under conditions that were not designed to capture transient or low-stoichiometry DNA methylation [1,4]. Yet it is now clear that conventional whole-genome bisulfite sequencing (WGBS) can introduce sequence-specific coverage and conversion biases that complicate interpretation at low abundance [4]. Enzymatic methyl-seq (EM-seq) and orthogonal chemistries have since improved sensitivity and fidelity in difficult settings, particularly at low input and in GC-rich or damage-prone regions [5–7]. In parallel, targeted HPLC/LC–MS pipelines coupled to authentic standards and NMR overlays provide definitive chemical identity for modified deoxynucleosides, enabling rigorous calls near the analytical limit of detection [16–19].

The *S. pombe* genome encodes Pmt1, a homolog of the Dnmt2/TRDMT1 family [9]. Dnmt2 proteins were long annotated as DNA methyltransferases by sequence homology, but seminal work showed that they methylate cytosine-38 in the anticodon loop of specific tRNAs, notably tRNA^Asp^, and protect tRNAs from stress-induced cleavage [10–13]. In *S. pombe*, Pmt1 methylates tRNA^Asp^ in vivo, with activity modulated by nutrient status via Sck2 signaling and by queuosine in the anticodon loop, which can stimulate Dnmt2 catalysis and alter tRNA decoding properties [9–11,14,15]. These observations raise a fundamental question: can a Dnmt2-family enzyme in a yeast widely considered “DNA-unmethylated” also catalyze DNA 5mC formation under specific physiological contexts?

Here we uncover a Pmt1-dependent DNA 5mC signal in *S. pombe*. Using validated nucleoside-level analytics (HPLC/LC–MS with external calibration and ICH Q2(R2)–compliant LOD/LOQ) and NMR confirmation of collected 5mdC fractions [16–20], we detect 5mdC that (i) rises acutely as cells re-enter the cell cycle from nitrogen-starvation quiescence (G0→S), (ii) is abolished in *pmt1Δ*, and (iii) accumulates in *fcy1Δ* backgrounds lacking the cytosine deaminase Fcy1, consistent with deamination-driven turnover of methyl-cytosine [21]. Live-cell Rad22-YFP imaging and DNA-content cytometry place this methylation pulse in the first S phase after quiescence, a window characterized by elevated recombination/repair foci and replication stress in *S. pombe* [22–26]. Together, these data identify Pmt1 as a bona fide DNA methyltransferase in fission yeast and suggest that 5mC formation is cell-state dependent, transient, and actively countered by base deamination. Beyond resolving a long-standing controversy for *S. pombe* [1–3], our work motivates a reevaluation of Dnmt2 catalytic plasticity and the contexts in which low-stoichiometry DNA methylation can be biologically meaningful.

## Methods

### Strains, media, and general conditions

Isogenic *Schizosaccharomyces pombe* strains were wild type, *pmt1Δ*, *fcy1Δ* and *fcy1Δpmt1Δ*. Cells were grown at 30 °C in YES or EMM with agitation (200 rpm) unless stated otherwise. The Rad22-YFP marker was introduced by standard genetics and expressed from its native locus.

### Quiescence (G0) induction and re-entry time course

Quiescence was induced by nitrogen starvation in EMM-N following established *S. pombe* workflows. Re-entry (“t = 0”) was triggered by transfer to nitrogen-replete medium. Samples were collected across the first cell cycle, with the first S-phase window annotated by DNA-content cytometry and septation index as in prior quiescence/exit frameworks [27, 44, 45]. Viability was monitored by colony-forming units on YES plates.

### Genomic DNA isolation and RNA removal

Cell pellets were lysed mechanically/enzymatically. Genomic DNA was purified using phenol-free buffers or silica-column kits, followed by sequential RNA depletion (RNase A/T1 → RNase H) and a short alkaline hydrolysis to eliminate residual RNA. DNA quality was assessed by A260/280 and A260/230 ratios, absence of rRNA bands on agarose gels, and dsDNA-specific fluorimetry. This workflow minimizes RNA carry-over in nucleoside assays [56].

### Enzymatic digestion to nucleosides

Defined masses of genomic DNA were digested to nucleosides using a standard sequential cocktail: DNase I (Mg²⁺, pH ≈ 7.0) → nuclease P1 (Zn²⁺, pH ≈ 5.2) → alkaline phosphatase (pH ≈ 7.5). Reactions proceeded to completion (verified by plateau of total nucleoside yield) and were filtered prior to chromatography. The DNase-I → NP1 → AP pipeline is widely adopted for quantitative nucleoside analysis and limits partial- hydrolysis bias [50, 49, 47].

### HPLC/LC–MS identification and quantification of dC and 5mdC

Digests and authentic standards (2′-deoxycytidine, 5-methyl-2′-deoxycytidine) were injected in the same analytical sessions. Separation used a reversed-phase column (C18, 3–5 µm) with aqueous/organic eluents compatible with UV (∼254 nm) and positive-mode electrospray. Identity of 5mdC in genomic digests was assigned by co-elution with the standard and MS signatures: [M+H] ⁺ = 242.1 in full scan and a (2M+H)⁺ ≈ 484.3 feature in product-ion spectra. External calibration curves converted areas to amounts; representative chromatograms and linear fits (R²) are provided in Supplementary Figure. S1 [16, 17, 18, 19, 47–49].

### Analytical validation, LOD/LOQ, and calling rules

Method validation followed ICH Q2(R2) (effective 2024). For 5mdC, the limit of detection (LOD) - calculated from calibration slope and baseline noise on blanks/low standards - was 0.0125 mM in our configuration; the limit of quantification (LOQ) was derived using ICH equations. “ND” (not detected) is defined as < LOD. Unless stated, values are mean ± SD from n ≥ 6 biological replicates. Inter- and intra-assay precision were assessed on pooled digests analyzed across days [20].

### NMR confirmation of 5mdC identity

HPLC fractions corresponding to putative 5mdC were collected and concentrated. ¹H (and, where available, **¹**³C) NMR spectra were acquired in D₂O and overlaid with a commercial 5mdC standard. Chemical-shift concordance across hallmark resonances (C5/H5, C6/H6, deoxyribose protons) and the absence of extraneous peaks supported molecular identity [16, 19, 56].

### Time-course and two-state snapshot designs

For the time course, aliquots were collected at fixed intervals after refeeding; for the two-state snapshot, samples were taken in vegetative growth and at **+**2 h after G0 exit. At each point, % 5mdC/ (5mdC + 5dC) was measured by HPLC/LC–MS as above. Wild type, *fcy1Δ*, *pmt1Δ*, and *fcy1Δ pmt1Δ* were assayed side-by-side within the same analytical sessions. ND calls strictly followed the LOD rule [27, 20].

### DNA-content flow cytometry

Cells were ethanol-fixed, treated with RNase A, stained with propidium iodide, and analyzed on a benchtop flow cytometer. G1/S/G2 distributions were used to delineate the first S-phase after G0 exit and to synchronize figure annotations with biochemical sampling [27, 44].

### Live-cell imaging of Rad22-YFP and focus quantification

Cells expressing Rad22-YFP were imaged on EMM pads with constant exposure/laser settings. Foci were scored blinded to genotype as nuclei containing ≥ 1 Rad22-YFP focus; ≥ 200 cells per condition per replicate were counted when feasible. Rad22-positive foci are established recombination/repair assemblies that increase under replication stress in *S. pombe*, providing a live-cell readout during G0→S [22, 23, 51].

### Queuine treatment during G0 exit and Rad22-YFP foci analysis

#### Strains and culture conditions

All strains carry a C-terminal Rad22-YFP fusion at the endogenous *rad22* locus. The four genotypes analyzed were wild type (wt), *pmt1Δ*, *fcy1Δ*, and *fcy1Δ pmt1Δ*. Cells were grown at 30 °C in EMM2 minimal medium with supplements to mid-log phase, then shifted to nitrogen-free EMM (EMM-N) to induce quiescence (G0) as described in the Methods (typically 24 h starvation unless stated).

### Nascent DNA preparation and sucrose-gradient fractionation (Okazaki-enrichment)

Cells were sampled at the indicated times after G0 release (Fig. 4A). Low-molecular-weight DNA was prepared and resolved on 10–40% sucrose gradients in nuclease-free buffer. Gradients were run on a piston-driven fractionator; 12 fractions (F1–F12) were collected at 50 µL each from top to bottom. Fractions were analysed on agarose gels to assess size distributions across the gradient (Fig. 4B). Where required, equal volumes for a given fraction index were pooled across time points (e.g., F1 = F1_G0 + F1_1h40 + …) to obtain sufficient material for enzymology and analytics. The workflow follows short-nascent-strand abundance/enrichment best practices [65,66].

### λ-Exonuclease enrichment of nascent strands

Pooled or individual fractions were incubated with λ-exonuclease under supplier-recommended conditions to digest 5′-phosphorylated dsDNA while sparing RNA-primed nascent strands that are refractory to 5′→3′ digestion [64,67]. Reactions were terminated, DNA recovered, and an aliquot run on agarose to confirm depletion of λ-exo–sensitive species. We adopted published precautions to limit known λ-exo sequence/structure biases (matched inputs, fraction-based pooling, no-enzyme controls) [64,65].

### HPLC quantification of 5mdC in nascent DNA

λ-exo–treated DNA from F1–F6 was enzymatically digested to nucleosides and analyzed by HPLC as in the main text (calibration/LOD in Fig. S1). %5mdC was computed as 5mdC/(5mdC+dC) × 100. Fractions from wt and *fcy1Δ* were processed side-by-side; biological replicates as indicated in Fig. 4D.

#### Release from G0 and queuine treatment

G0 cultures were harvested, washed once in pre-warmed EMM2, and released into fresh EMM2 (time 0). Immediately upon release, cells were split into ±queuine conditions and incubated for 1 h at 30 °C with shaking.

#### Queuine (+Q)

queuine base (hemi-sulfate), stock 10 mM in sterile water, filter-sterilized, added to a final concentration of 10 µM. Unless indicated, 10 µM was used for all experiments. After 1 h, samples were processed for imaging (below). In parallel, the same ±Q protocol was applied to each genotype (wt, *pmt1Δ*, *fcy1Δ*, *fcy1Δ pmt1Δ*).

#### Statistics

Effects of genotype and treatment (±Q) were tested by two-way ANOVA (Prism software). Significance thresholds were pre-specified (e.g., *p* < 0.05, 0.01, 0.005 as annotated in figure legends). Graphs display mean ± SD; exact *n* and *p* values are provided in the legends (see Fig. S2).

## Results

To determine whether fission yeast carries a minimal yet functional DNA methylation program, we analyzed quiescent cells as they re-entered the cell cycle and combined three layers of evidence: physiology (cell-cycle progression and DNA-repair foci), direct nucleoside analytics, and genetics. This integrative design avoids conversion-dependent biases and enables quantitative, state-resolved measurements of 5-methyl-2′-deoxycytidine (5mdC) in genomic DNA [16–19,20].

We first characterized the re-entry trajectory from G0. DNA-content cytometry defined the timing of the first S phase after refeeding, and revealed a modest G1/S delay in the cytosine/5-mC deaminase mutant *fcy1Δ* relative to wild type (wt) (Figure. 1A). Live-cell imaging of Rad22-YFP - an established marker of recombination-repair assemblies - showed few or no foci in quiescence but a pronounced increase during this first S phase; importantly, *fcy1Δ* accumulated a higher fraction of nuclei with Rad22 foci than wt at matched time points (Figure. 1B–C) [22–26]. These observations place the forthcoming chemistry into a replication-linked context and motivate a search for methyl-cytosine dynamics during G0→S.

**Figure 1:**
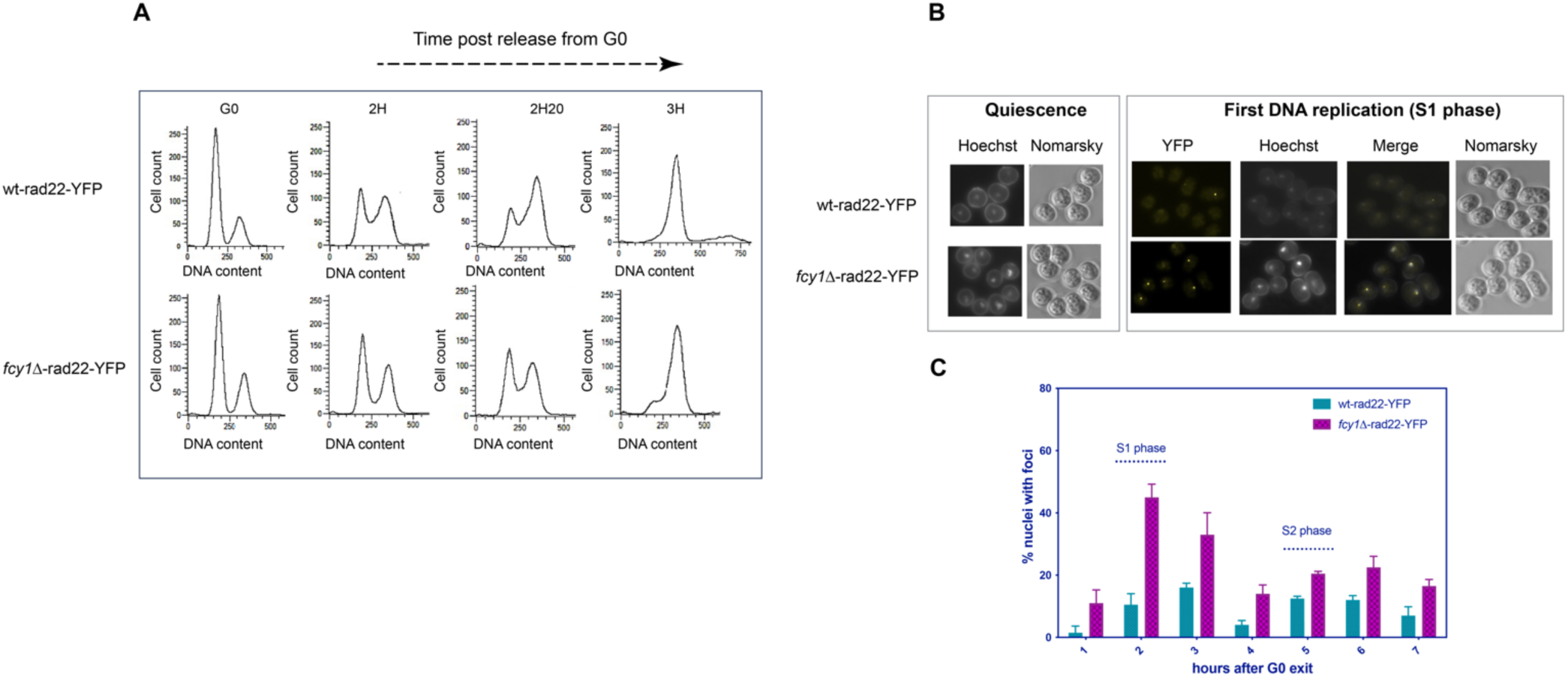
Loss of cytosine deaminase *(fcy1Δ)* delays G1/S after GO exit and increases spontaneous Rad22-YFP DNA-repair foci. **A:** DNA-content profiles (FAGS) of wt-rad22-YFP and fcy1Δ-rad22-YFP during re-entry from GO highlight a G1/S delay in *fcy1Δ.* **B:** Representative micrographs showing spontaneous Rad22-YFP foci in quiescent cells and during the first S phase. **C:** Quantification of Rad22-YFP-positive nuclei over time (mean± SD, n ≥ 3 biological replicates; ≥ 200 nuclei per condition per replicate). Statistics: two-way ANOVA (genotype x time) with multiple-comparisons correction; **** p < 0.0001. These data indicate replication-associated genome stress when methyl-cytosine turnover is impaired.

We next asked whether genomic 5mdC can be detected and identified unambiguously. Enzymatic digestion of genomic DNA to nucleosides followed by HPLC/LC–MS revealed a peak co-eluting with an authentic 5mdC standard; full-scan spectra displayed the diagnostic (M+H)^+^ ion at m/z 242.1 and product-ion spectra showed the expected (2M+H)^+^ feature at ∼484.3 (Figure. 2A–C). External calibrations for dC and 5mdC established linearity and sensitivity (limit of detection, LOD = 0.0125 mM), supporting quantitative calls including “ND” (Figure. S1); this workflow follows ICH Q2(R2) - style validation and circumvents conversion chemistry artefacts [16–20].

**Figure. 2.**
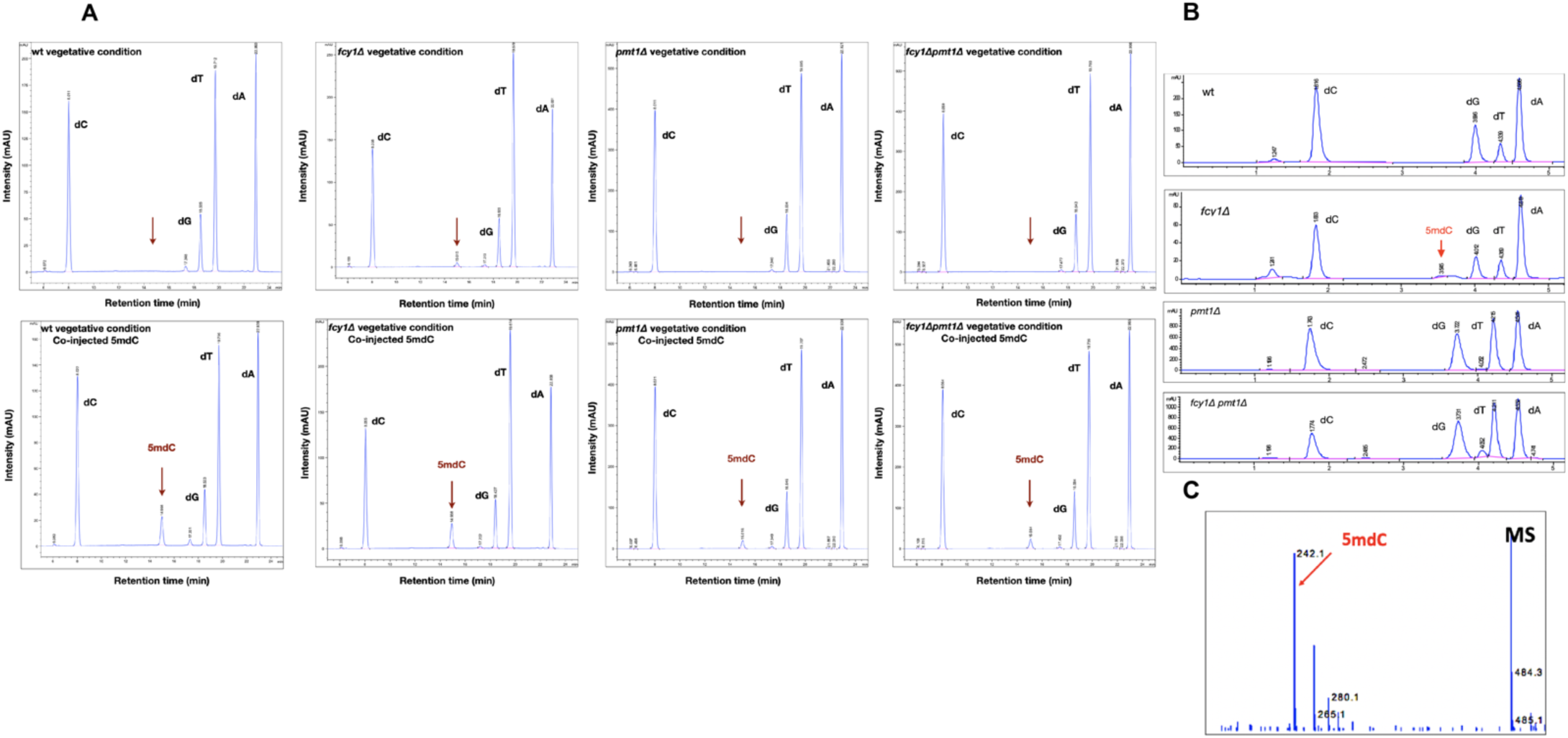
Chemical detection and identity of genomic 5-methyl-2’-deoxycytidine (5mdC) in S. *pombe.* on the Pmt1 DNA methyltransferase (HPLC/LC-MS). **A:** HPLC chromatograms of enzymatic DNA digests from wild-type (wt), fcy1Δ pmt1Δ and fcy1Δ, pmt1Δ, cells in vegetative conditions. Co-injection of an authentic 5mdC standard demonstrates co-elution (red arrows). **B:** LC-MS full-scan spectra for the putative 5mdC peak show the diagnostic [M+H]+ ion at m/z 242.1. **C** Product-ion spectrum reveals the (2M+H)+ adduct at ∼m/z 484.3, confirming molecular identity. Together, orthogonal chromatographic and mass-spectrometric signatures establish authentic 5mdC in genomic digests.

Using this validated assay, we then quantified 5mdC across cell state and genotype. In wt, 5mdC was undetectable in vegetative growth but rose transiently during the first S phase after G0 exit (Figure. 3A–C). Deletion of the Dnmt2-family methyltransferase Pmt1 abolished the signal at all time points, demonstrating that the DNA-embedded 5mC we detect is Pmt1-dependent. Conversely, loss of Fcy1 markedly amplified the same S-phase-linked pulse and elevated residual 5mdC at steady state, consistent with deamination-driven turnover limiting methyl-cytosines as cells re-enter the cycle (Figure. 3A–C) [3–7,21]. As a representative snapshot, wt increased from ND (vegetative) to ∼0.52% ± 0.03 at +2 h, whereas *fcy1Δ* rose from 1.45% ± 0.36 to 5.94% ± 1.36 at +2 h; *pmt1Δ* and *fcy1Δ pmt1Δ* remained ND (means ± SD; n ≥ 6 biological replicates).

**Figure 3.**
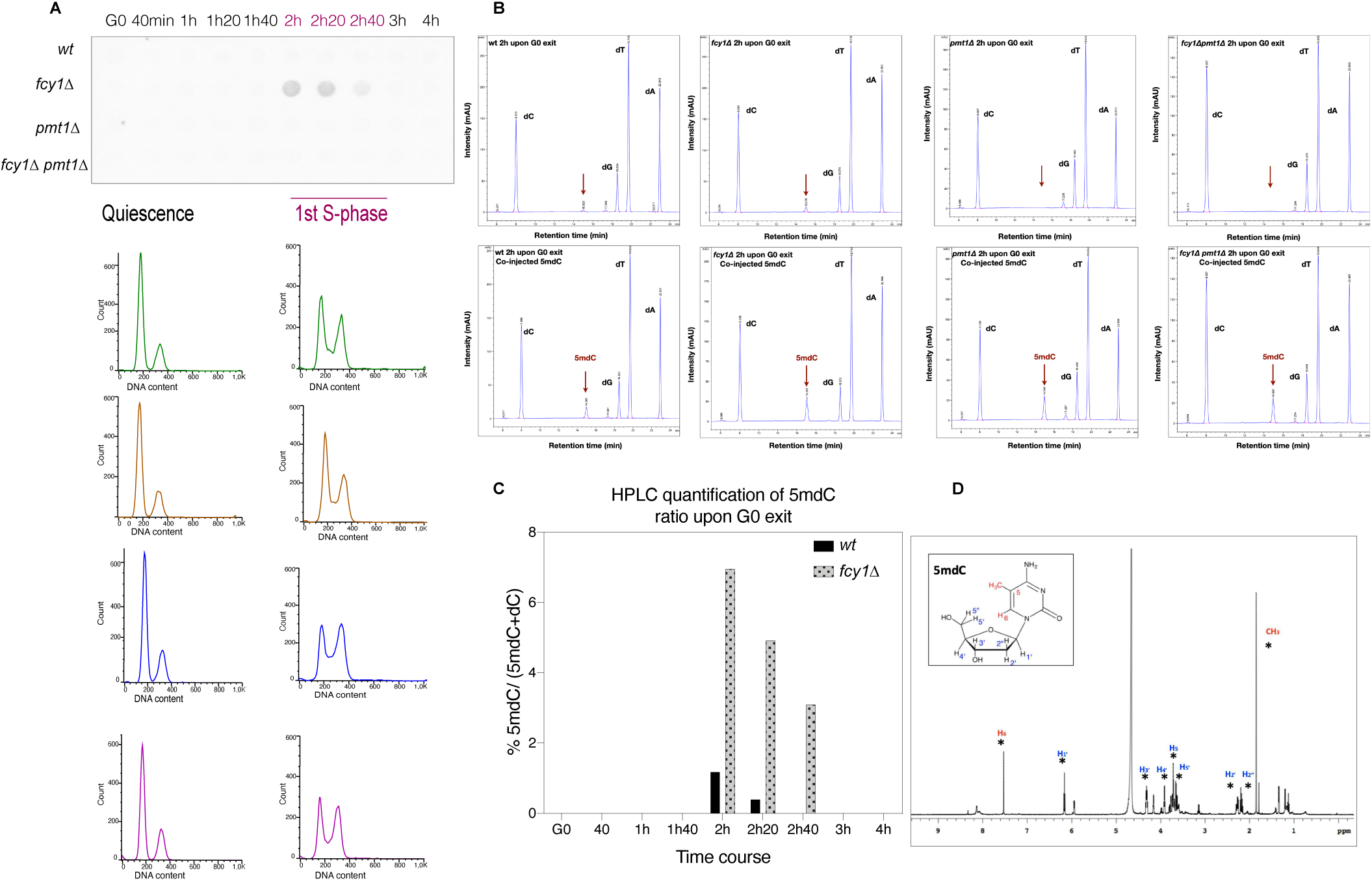
5mdC transiently rises at the first S phase upon GO exit and is modulated by Pmt1 and Fcy1; NMR confirmation. **A:** Dot-blot of methylated DNA (anti-5mC) across the GO-exit time course (top) and DNA-content profiles (FACS) demarcating quiescence and the first S phase (bottom) for wt, *fcy1Δ, pmt1Δ and fcy1Δ pmt1Δ.* **B:** HPLC/LC-MS chromatograms at +2 h after GO exit for each genotype; co-injected 5mdC standard (bottom panels) confirms co-elution (red arrows). **C:** Quantification of %5mdC/(5mdC+dC) across time shows a transient pulse in wt that is abolished in *pmt1Δ.* and strongly amplified in *fcy1Δ..* **D:** 1H-NMR spectrum of the HPLC-collected 5mdC fraction *(fcy1Δ.)* overlays the commercial standard, validating chemical identity.

**Figure 4.**
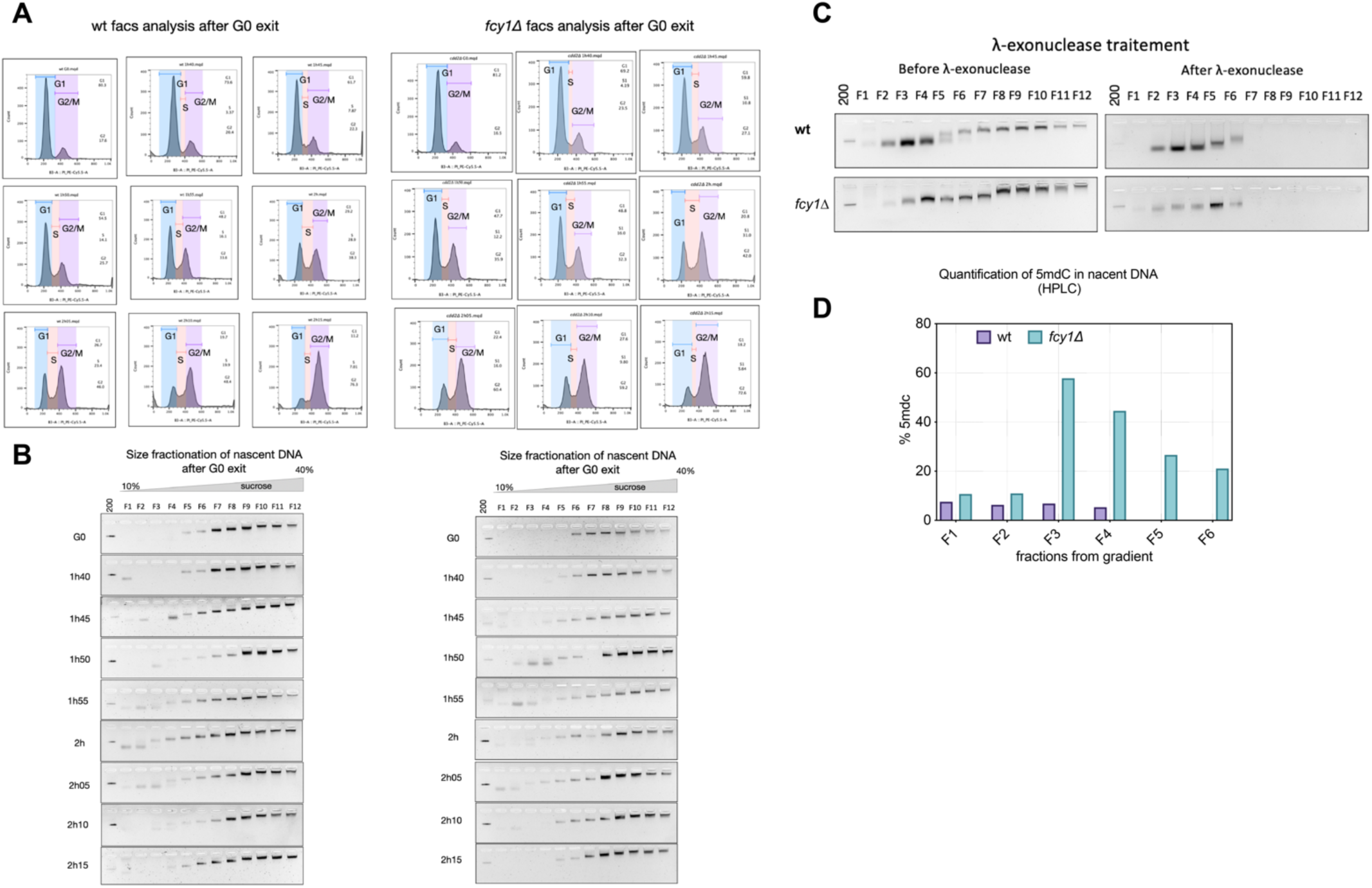
Nascent (Okazaki-rich) DNA carries the Go→s SmdC pulse. **A.** FAGS profiles during GO exit (wt vs *Fcy1Δ, named cdd2Δ at the begain of study)* to stage S phase. **B.** Size-fractionation of nascent DNA on 10-40% sucrose gradients, wt and *Fcy1Δ* (12 fractions, 50 µLeach) at the indicated times. **C.**λ A-exonuclease test on pools by fraction index (e.g., all F1 from every time point mixed): signal resistant after λ-exo marks NA-primed nascent strands (Okazaki-enriched). **D.** HPLC of F1-F6 after λ-exo: %5mdC peaks in mid-fractions and is elevated in *Fcy1Δ* versus wt, consistent with a Pmt1-installed, Fcy1-restrained S-phase methylation pulse.

To further secure chemical identity, the HPLC-collected peak was analyzed by 1H-NMR, which overlaid the commercial 5mdC standard across hallmark resonances (Figure. 3D). Taken together with Figure. 1, these data define a Pmt1↔Fcy1 axis that installs and drains a short-lived 5mdC pulse precisely at G0→S in Schizosaccharomyces pombe [3–7,16–21].

To position fcy1 within the quiescent mutagenesis network, we leveraged the sensitized BER/THO background *ung1Δ thp1Δ*, in which our recent work showed enhanced mutation accumulation during G0 [63]. Across the G0 time course, *ung1Δ thp1Δ* displayed reduced viability relative to wt and a clear rise in FOAR frequency (Figure. S3A–B). The forward-mutation spectra were most informative: at day 1, *ung1Δ thp1Δ* was dominated by C→T (G→A) transitions, a hallmark of uracil-type lesions that are normally excised by Ung1; this transition bias persisted at day 16 (Figure. S3C–D). Critically, adding *fcy1Δ* on top of *ung1Δ thp1Δ fcy1Δ* attenuated FOAR accumulation and shifted the spectra away from transitions, with an ≈30% drop in C→T at day 16 relative to the double mutant (Figure. S3B–D). Together with the Pmt1-dependent 5mdC pulse at G0→S (Figure. 3A–C) and the Rad22-YFP phenotypes (Figure. 1), these data indicate that Fcy1- mediated deamination (of C and/or 5mC) contributes materially to the mutagenic lesion burden in quiescence - especially when Ung1-dependent uracil excision and Thp1/THO-dependent RNA: DNA hybrid control are compromised.

To determine whether the transient 5mdC detected after G0 exit is associated with newly synthesized strands, we isolated nascent DNA by sucrose-gradient fractionation and λ-exonuclease enrichment, following best practices from nascent-strand (NS-seq) workflows (Gerbi and colleagues) with some adaptations to *S. pombe* (Figure. 4A–C) [64–66]. Fractions corresponding to the Okazaki-size window retained signal after λ- exo - consistent with protection by 5′ RNA primers - whereas larger/parental fragments were largely removed (Figure. 4C) [64,67]. HPLC on the λ-exo–enriched material revealed 5mdC above LOD in F1–F6, with a peak in mid-fractions (≈F3–F5) and a marked elevation in *fcy1Δ* relative to wt (Figure. 4D). These data place the Pmt1-dependent 5mdC pulse on nascent DNA strands, consistent with lagging-strand (Okazaki) - rich material, and strengthen the view that Fcy1 limits the methyl-cytosine pool immediately after DNA synthesis [64–66,68]. We note λ-exo biases and controlled for them by fraction-matched pooling and side-by-side genotype controls (Methods) [64,65].

Finally, because Dnmt2/Pmt1 activity is known to respond to nucleotide and queuosine/queuine metabolism in RNA contexts [9–15], we tested whether queuine modulates the replication-linked phenotype. Supplementation reduced Rad22-YFP foci in *fcy1Δ* during G0 exit (Figure. S2), suggesting that metabolic inputs can tune this methylation–deamination axis during cell-cycle re-entry.

## Discussion

Our work overturns the long-standing view that the fission yeast *Schizosaccharomyces pombe* lacks genomic DNA 5-methylcytosine (5mC) [1–3]. Using orthogonal chemistry - HPLC/LC–MS with external calibration and ICH-compliant sensitivity, together with NMR overlays of collected fractions - we unambiguously identify 5-methyl-2′-deoxycytidine (5mdC) in genomic DNA digests [16–20]. Genetic dissection places this modification squarely on the Dnmt2-family enzyme Pmt1: 5mC is abolished in *pmt1Δ* and markedly elevated in *fcy1Δ*, which lacks the cytosine/5mC deaminase Fcy1 [21]. Temporally, the signal peaks at the first S phase as cells exit quiescence (G0→S), and enlarged methyl-cytosine pools in *fcy1Δ* coincide with Rad22- YFP foci diagnostic of replication-coupled recombination [22–26]. Together, these data establish Pmt1 as a bona fide DNA methyltransferase in vivo and reveal an actively regulated Pmt1↔Fcy1 axis that restricts methyl-cytosine during cell-cycle re-entry.

### Reconciling prior “no 5mC” reports with a transient, regulated methylome

Earlier surveys concluded that *S. pombe* is essentially unmethylated [1–3], but those studies were not optimized to capture low-stoichiometry, cell-state-restricted DNA methylation. Bisulfite library preparation can under-represent difficult contexts and inflate apparent variation through conversion and coverage biases, especially when true methylation is scarce [4]. Enzymatic approaches and direct chemistry have since raised the bar for trace-level detection [5–7,16–20]. Three features of our dataset help explain the discrepancy: (i) temporal restriction - a short 5mC pulse at G0→S is diluted in asynchronous populations; (ii) analytical rigor - a validated LOD (0.0125 mM) and run-matched standards make “ND” calls meaningful; and (ii) genetic anchoring - side-by-side *pmt1Δ/fcy1Δ* comparisons expose causal control.

Our genetics separate installation and turnover: Pmt1 (Dnmt2) installs DNA 5mC in vivo, while Fcy1 drains 5mC/C through deamination its loss amplifies and prolongs the G0→S 5mC pulse and coincides with replication-linked Rad22 foci (Figures. 3, 1) [3–7,21,22–26]. Epistasis in a sensitized background sharpens this model: in *ung1Δthp1Δ*, which accumulates uracil-type C→T transitions in G0, removing *fcy1* (triple mutant *fcy1Δung1Δthp1Δ*) reduces FOA^R^ accumulation and lowers C→T by ∼30% (Figure. S3), placing Fcy1-mediated deamination upstream as a proximal source of mutagenic uracil/thymine lesions when uracil excision (Ung1) and R-loop control (Thp1/THO) are compromised [63]. In wt, the Pmt1↔Fcy1 balance keeps the 5mC/C deamination flux minimal and transient - installing a short signal at G0→S then rapidly curtailing it - thereby limiting T: G/U: G mismatches and the engagement of error-prone repair.

### Origin-proximal, co-replicative installation of 5mdC

Locating the 5mdC pulse on λ-exo–enriched nascent DNA argues that Pmt1 acts co-replicatively, most conspicuously within the Okazaki fragment size range (Fig. 4) [64–66,68]. This fits the temporal peak at G0→S (Figure. 3) and the Rad22-YFP phenotypes (Figure. 1), and suggests that the Pmt1↔Fcy1 axis tunes methyl-cytosine exposure at or just behind replication forks. Given documented λ-exo biases in nascent-strand assays, future bisulfite-independent, strand-resolved mapping integrated with Okazaki-fragment sequencing will be informative, particularly to distinguish leading vs lagging strand deposition and to test for origin proximity [64,65,68]

### Functional rationale for a minimal, timed 5mC program

Fission yeast appears to practice epigenetic minimalism: DNA 5mC is deployed briefly and state-dependently as a pulse at G0→S rather than as a stable bulk mark. Such low-stoichiometry, regulated 5mC could fine-tune replication-fork transactions (e.g., origin usage or Okazaki-fragment maturation), temper the handling of repetitive DNA, and interface with stress pathways during nutritional transitions. This functional sufficiency at low abundance aligns with the heterogeneity of fungal methylomes and with the persistence of sparse methylation in lineages where canonical DNMTs have been lost or repurposed [3,41]. The broader Dnmt2/TRDMT1 literature also argues that catalytic environment and cofactors shape substrate choice: queuosine and nutrient signaling modulate Pmt1/Dnmt2 on tRNA [9–15], and S-adenosylmethionine availability shifts across quiescence and re-entry [45]. We therefore propose that cell state, chromatin access, and cofactor supply bias Pmt1 toward DNA transiently at G0→S, with Fcy1 rapidly curtailing the footprint - yielding a minimal, timed 5mC signal that is effective without being pervasive.

### Broader implications and DNMT2 dual-substrate capacity

Our findings in fission yeast are consistent with a broader view in which DNMT2/TRDMT1 enzymes can shape both RNA and DNA cytosine methylation depending on cellular context. In the malaria parasite Plasmodium falciparum, the global DNA cytosine modification landscape (5mC and oxidized derivatives) becomes DNMT2-dependent under controlled oxygen regimes, supporting a direct DNMT2-linked DNA methylation route in vivo [61]. Complementarily, DNMT2-mediated tRNA C38 methylation modulates stress tolerance and proteome allocation in Plasmodium, highlighting DNMT2’s context-dependent, dual-substrate biology [62]. Together with the Pmt1↔Fcy1 axis uncovered here, these observations support a model in which Dnmt2-family enzymes install a minimal, state-restricted DNA 5mC signal, while deamination pathways and metabolic inputs such as queuine constrain its magnitude and timing at key physiological transitions.

### Limitations and testable predictions

We have not yet localized sites genome-wide. Bisulfite-free mapping (for example EM-seq) should resolve sequence context and reveal whether Pmt1 favors particular CpN motifs or origin-proximal regions [5–7]. If the model is correct, catalytic-site mutants of Pmt1 (motif IV cysteine; motif I SAM binding) will abolish the pulse; in vitro assays with purified Pmt1 should methylate dsDNA substrates under conditions that mimic G0→S. The turnover model predicts epistasis with base-excision and mismatch repair, and that elevating Pmt1 activity outside G0→S will be toxic unless Fcy1 capacity or downstream processing increases. Finally, perturbing SAM metabolism or queuine supply should modulate pulse amplitude and timing, linking metabolic state to Pmt1 substrate choice [9–15,45].

### Implications beyond fission yeast

That a Dnmt2-family enzyme can methylate DNA in vivo in a eukaryote long considered essentially unmethylated broadens the functional landscape of this enzyme class [9–15]. Rather than a binary “RNA-only” versus “DNA-only” assignment, our results support a conditional, dual-substrate view in which cell state and counter-enzymes (here, Fcy1) determine when DNA methylation is deployed sparingly yet purposefully. *Schizosaccharomyces pombe* offers a genetically clean chassis to dissect how minimal methylomes are produced, sensed, and erased, and how this economy of marking influences genome maintenance during environmental transitions.

In summary, Pmt1 catalyzes DNA 5mC in *S. pombe*; the mark peaks transiently at G0→S, is eliminated in *pmt1Δ*, and is amplified in *fcy1Δ*, where it coincides with replication-coupled recombination foci. This combined chemical, genetic and temporal logic revises the epigenetic landscape of fission yeast and motivates bisulfite-independent mapping and mechanistic tests to define where Pmt1 methylates and how Fcy1 restrains this footprint.

These findings also align with emerging evidence outside fission yeast. In *Plasmodium falciparum*, the global DNA cytosine-modification landscape (5mC and oxidized derivatives) becomes DNMT2-dependent under defined oxygen regimes, consistent with a direct DNMT2-linked DNA methylation route in vivo [61]. Complementarily, DNMT2-mediated tRNA C38 methylation modulates stress tolerance and proteome allocation in *Plasmodium* [62]. Together with the Pmt1↔Fcy1 axis described here, these observations raise the possibility that other Dnmt2 enzymes exhibit cryptic, state-restricted DNA methylation under specific physiological conditions, with deployment gated by metabolic inputs and opposing deamination pathways.

### Data Origin, Figures, and Materials Availability

The data presented in this manuscript were generated as part of an independent research initiative that I conceived and developed during my postdoctoral training on 2016, in the laboratory of Dr. Benoît Arcangioli, with support from the *Fondation de France* (Prix Thérèse Lebrasseur). This work reports the first detection of DNA 5-methylcytosine (5mC) in *Schizosaccharomyces pombe* and identifies Pmt1 as the enzyme responsible for this modification. By depositing this work as a preprint on bioRxiv, I aim to ensure the permanent accessibility of these findings to the scientific community and to establish an official, traceable record of authorship.

All data, figures, and interpretations presented in this preprint are the intellectual property of the author and collaborators. This work is released under the Creative Commons Attribution (CC BY 4.0) license selected during submission. Any reproduction, reuse, or derivative work - including figures, datasets, or textual excerpts - must properly cite this preprint using its DOI and full bibliographic reference.

## Acknowledgments

I would like to express my deepest gratitude to Valérie Huteau and Dr. Sylvie Pochet *(Unité de Chimie et Biocatalyse, Institut Pasteur)* for their exceptional technical expertise, insightful discussions, and invaluable support throughout this study.

Their contributions were instrumental in the successful completion of the LC/MS validation experiments, which form a critical foundation of this work.

I also warmly thank Dr. Benoît Arcangioli *(Unite Dynamique du Génome, Institut Pasteur)* for his scientific mentorship and continuous guidance throughout the development of this project.

---

**Figure 1. *fcy1Δ* induces a G1/S delay upon G0 exit and accumulates spontaneous Rad22-YFP repair foci.**

(A) DNA-content profiles (FACS) at G0, +2 h, +2 h20, +3 h (wt-rad22-YFP; f*cy1Δ*-rad22-YFP).

(B) Representative micrographs in quiescence and during the first S phase (YFP, Hoechst, merge, Nomarski).

(C) Quantification of nuclei with Rad22-YFP foci across time after G0 exit; S-phase windows indicated. n>200 nuclei/strain; 3 biological replicates; two-way ANOVA, p<0.001.

**Figure 2.** Global genomic 5-methyl-2′-deoxycytidine (5mdC) is detectable and requires Pmt1 (HPLC/LC–MS).

(A) HPLC chromatograms of enzymatic DNA digests from wt, *fcy1Δ*, *pmt1Δ*, *fcy1Δ pmt1Δ* (top: native; bottom: + co-injected 5mdC). A co-eluting peak appears in wt and *fcy1Δ*, but not in *pmt1Δ* backgrounds.

(B) Full-scan LC–MS shows the protonated adduct [M+H]^+^ at m/z 242.1.

(C) Product-ion spectrum confirms 5mdC with a (2M+H)^+^ feature at ∼m/z 484.3.

**Figure 3.** 5mdC rises during the first S phase after G0 exit in *fcy1Δ* in a Pmt1-dependent manner.

(A) Dot-blot of genomic DNA probed for 5mC over a G0-exit time course.

(B) HPLC traces across time (wt vs fcy1Δ; + co-injected 5mdC).

(C) HPLC quantification of %5mdC/(5mdC+dC) upon G0 exit (mean±SD).

(D) 1H-NMR of the purified 5mdC fraction, matching an authentic standard.

## Supplementary Figure Legends

**Figure S1. Analytical validation of 5mdC/dC measurements.**

(A–B) External calibration curves for 5mdC and dC (linear fits, R2>0.99; LOD ∼0.0125 mM for 5mdC).

(C) HPLC chromatograms of standard nucleosides.

**Figure S2. Pmt1-dependent DNA damage upon G0 exit in f*cy1Δ* and suppression by queuine.**

(A) % nuclei with Rad22-YFP foci over time in wt, *fcy1Δ*, *pmt1Δ*, *fcy1Δ pmt1Δ*.

(B) Queuine (Q) reduces Rad22-YFP foci in *fcy1Δ*; n>250 nuclei/strain; 2 experiments; two-way ANOVA, p<0.005.

**Supplementary Figure S3. Quiescent mutagenesis in *ung1Δ thp1Δ* is dominated by uracil-type transitions and partially suppressed by *fcy1Δ*.**

**(A) Viability curves** during quiescence (G0) for *wt*, *fcy1Δ*, *ung1Δ thp1Δ*, and *fcy1Δ ung1Δ thp1Δ* strains. The triple mutant exhibits reduced survival compared to *wt*, while loss of *fcy1* partially rescues viability defects in the double mutant background.

**(B) Accumulation of FOA-resistant (FOA^R^) colonies** over time in G0. The *ung1Δ thp1Δ* double mutant displays a strong increase in forward mutation frequency, which is attenuated in the *fcy1Δ ung1Δ thp1Δ* background, indicating that uracil removal by Ung1 contributes substantially to mutation burden.

**(C–D) Forward-mutation spectra** at day 1 (**C**) and day 16 (**D**) in G0. Mutation profiles reveal a predominance of C→T / G→A transitions in *ung1Δ thp1Δ*, consistent with unrepaired uracil lesions generated by cytosine deamination or misincorporation. The *fcy1Δ* deletion partially suppresses these transitions, confirming its role in limiting methyl-cytosine–derived uracil accumulation.

**Figure 4.** Nascent DNA (Okazaki fragments) carries the G0→S 5mdC pulse: sucrose-gradient fractionation, λ-exonuclease enrichment and HPLC quantification.

**(A)** Cell-cycle staging across the G0-exit time course by DNA-content cytometry (wt vs *fcy1Δ*). **(B)** Size-fractionation of nascent DNA on 10–40% sucrose gradients at the indicated times (G0 → +2 h 15); 12 fractions (F1–F12) were collected per gradient (50 µL/fraction) using the same piston system employed for ribosome gradients. **(C)** λ-exonuclease test of pooled fractions: for each fraction index (F1…F12), equal volumes from all-time points were combined (e.g., F1_G0 + F1_1h40 + …) to increase material, then analyzed before and after λ-exo. Loss of the bulk signal after λ-exo indicates depletion of broken/parental DNA carrying 5′-phosphate ends, whereas RNA-primed nascent strands remain enriched [64–67]. **(D)** HPLC quantification of %5mdC in F1–F6 after λ-exo (wt vs *fcy1Δ*). 5mdC is detectable and enriched in mid-fractions (Okazaki-size window) and is substantially higher in *fcy1Δ* than wt, consistent with a Pmt1-installed, Fcy1-restrained methylation pulse during S phase. Bars show means; exact *n*, enzyme units and gradient parameters are in Methods. We note the known sequence/structure biases of λ-exo selections and mitigated them as in [64,65].

## Supplementary Figures

**Figure S1:**
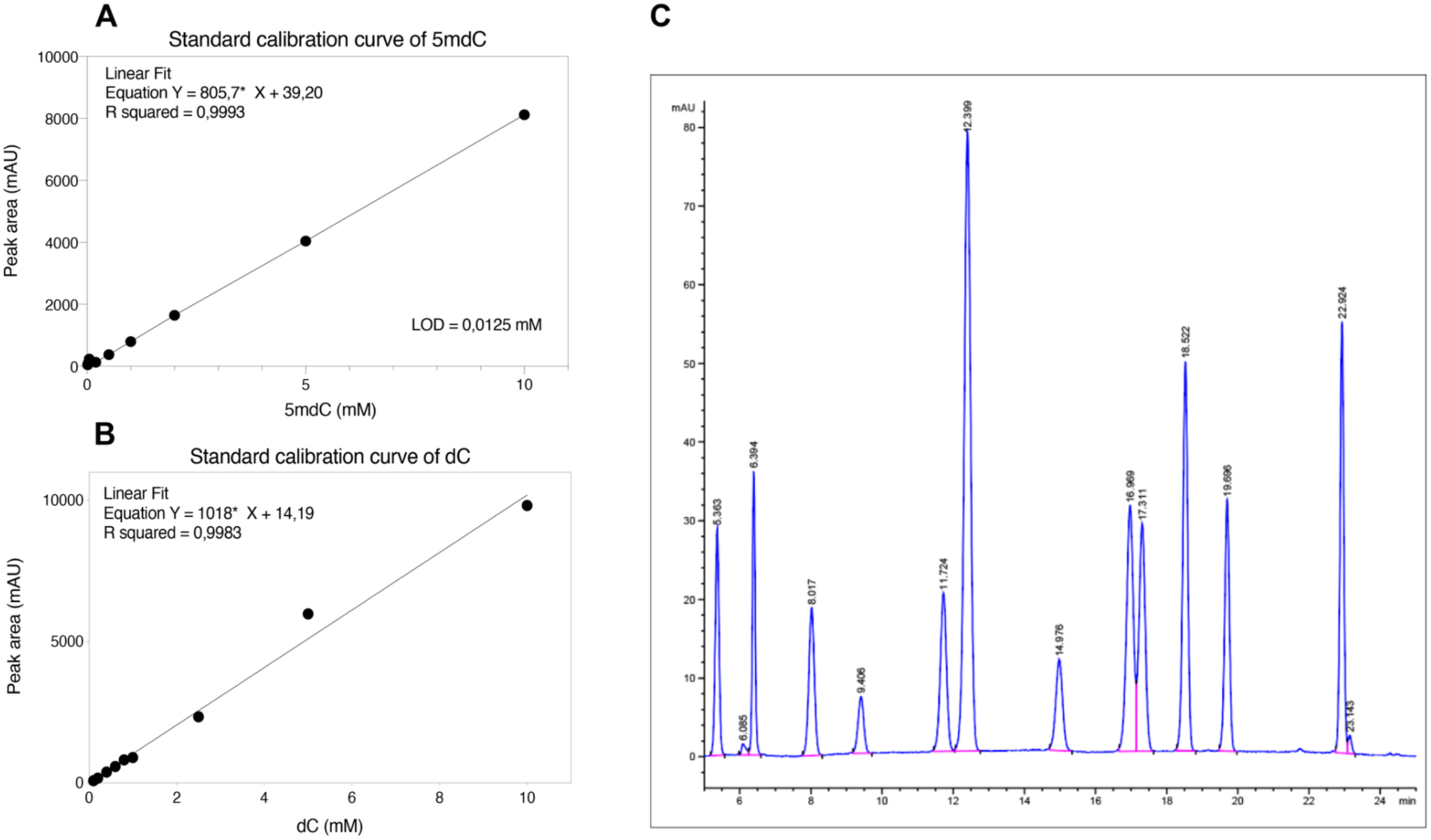
Calibration and standards for nucleoside analytics. **A:** External calibration curve for 5-methyl-2’-deoxycytidine (5mdC); linear fit shown with equation and R^2^. **B:** Calibration curve for 2’-deoxycytidine (dC). **C:** HPLC chromatogram of standard nucleosides used to identify and quantify dC and 5mdC. The limit of detection (LOO) for 5mdC in our setup is 0.0125 mM.

**Figure S2:**
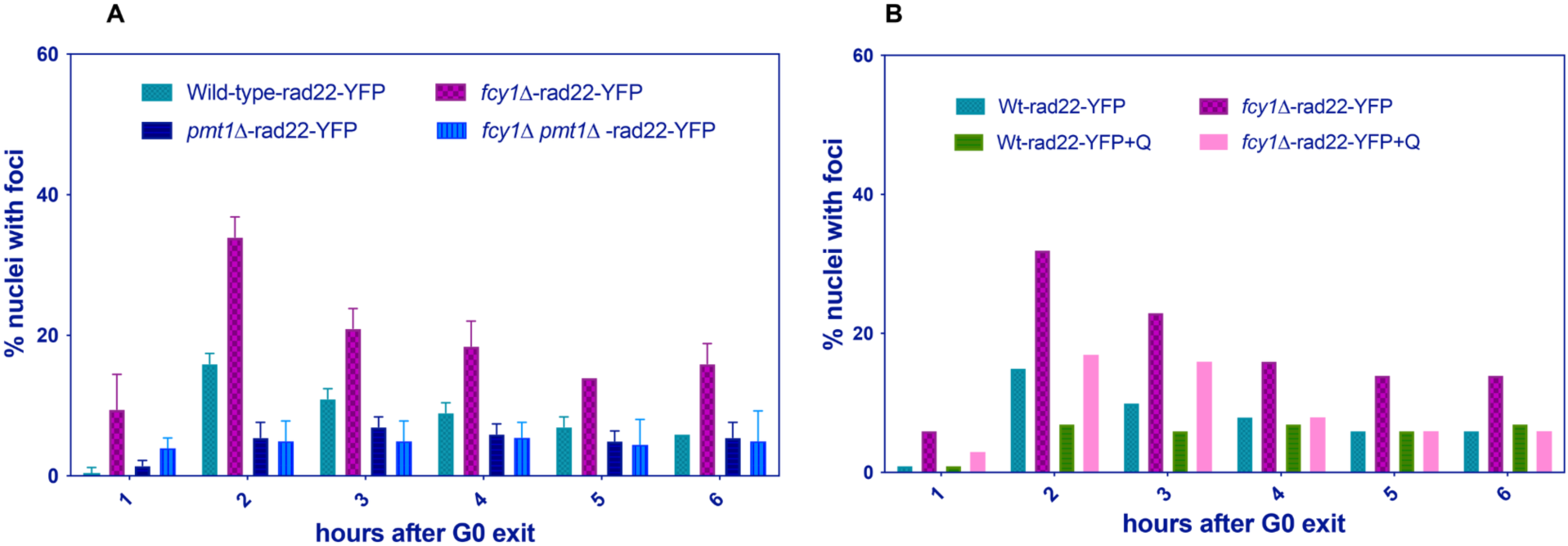
Rad22-YFP foci after GO exit across genotypes and modulation by queuine. **A:** Percentage of nuclei with Rad22-YFP foci over time in wt-, *fcy1Δ-, pmt1Δ-* and *fcy1Δ* pmt1Δ-rad22-YFP strains, (mean± SD; > 250 nuclei per strain). **B:** Effect of queuine supplementation (+Q) on Rad22-YFP foci in wt and *fcy1Δ* backgrounds during GO exit. Statistics: two-way ANOVA with appropriate post-hoc tests; p < 0.005 where indicated.

**Figure S3.**
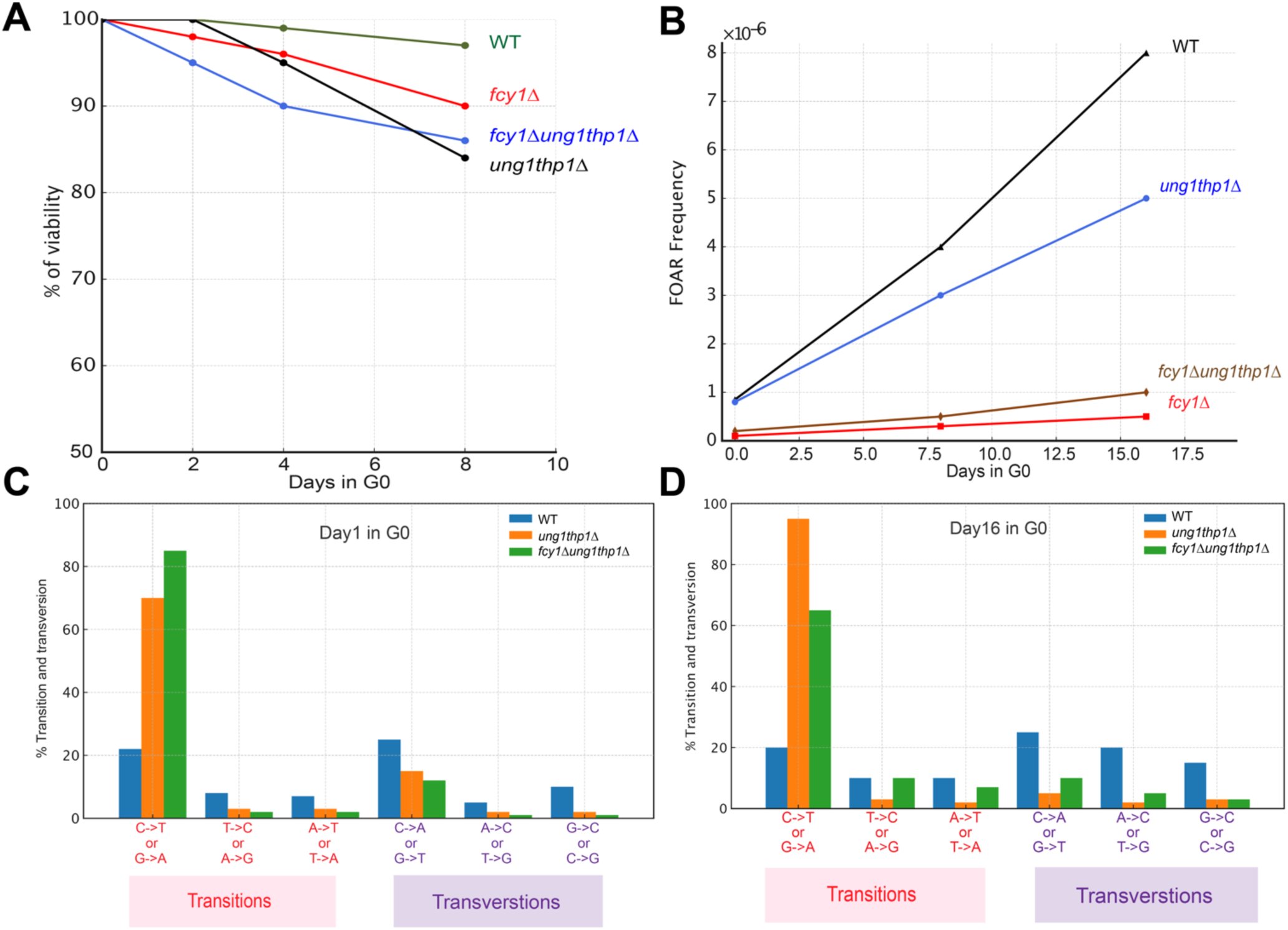
Quiescent mutagenesis in *ung111 thp111* is dominated by uracil-type transitions and is partially suppressed by *fcy1Δ.* **(A)** Viability during quiescence (GO) for wt, *fcy1Δ, ung1Δ thp1Δ,* and *fcy1Δ ung1Δ thp1Δ.* **(B)** Accumulation of 5-FOA-resistant (FOAR) colonies over the same time course. **(C-D)** Forward-mutation spectra at day 1 and day 16 in GO

